# Video-Based Deep Learning to Detect Dyssynergic Defecation with 3D High-Definition Anorectal Manometry

**DOI:** 10.1101/2021.12.11.472233

**Authors:** Joshua J. Levy, Christopher M. Navas, Joan A. Chandra, Brock C. Christensen, Louis J. Vaickus, Michael Curley, William D. Chey, Jason R. Baker, Eric D. Shah

**Affiliations:** Section of Gastroenterology and Hepatology, Dartmouth-Hitchcock Health, Lebanon, NH, USA; Quantitative Biomedical Sciences, Geisel School of Medicine at Dartmouth, Lebanon, NH, USA; Department of Epidemiology, Geisel School of Medicine at Dartmouth, Lebanon, NH, USA; Departments of Pharmacology and Toxicology, and Community and Family Medicine, Geisel School of Medicine at Dartmouth, Lebanon, NH, USA; Emerging Diagnostic and Investigative Technologies, Department of Pathology and Laboratory Medicine, Dartmouth-Hitchcock Health, Lebanon, NH, USA; Division of Gastroenterology and Hepatology, Michigan Medicine, Ann Arbor, MI, USA; Atrium Motility Laboratory, Division of Gastroenterology, Atrium Health, Charlotte, NC, USA

**Keywords:** artificial intelligence, machine learning, gastrointestinal motility, rectal diseases, Diagnostic Tests, Routine

## Abstract

**BACKGROUND AND AIMS:** Evaluation for dyssynergia is the most common reason that gastroenterologists refer patients for anorectal manometry, because dyssynergia is amenable to biofeedback by physical therapists. High-definition anorectal manometry (3D-HDAM) is a promising technology to evaluate anorectal physiology, but adoption remains limited by its sheer complexity. We developed a 3D-HDAM deep learning algorithm to evaluate for dyssynergia.

**METHODS:** Spatial-temporal data were extracted from consecutive 3D-HDAM studies performed between 2018-2020 at a tertiary institution. The technical procedure and gold standard definition of dyssynergia were based on the London consensus, adapted to the needs of 3D-HDAM technology. Three machine learning models were generated: (1) traditional machine learning informed by conventional anorectal function metrics, (2) deep learning, and (3) a hybrid approach. Diagnostic accuracy was evaluated using bootstrap sampling to calculate area-under-the-curve (AUC). To evaluate overfitting, models were validated by adding 502 simulated defecation maneuvers with diagnostic ambiguity.

**RESULTS:** 302 3D-HDAM studies representing 1,208 simulated defecation maneuvers were included (average age 55.2 years; 80.5% women). The deep learning model had comparable diagnostic accuracy (AUC=0.91 [95% confidence interval 0.89-0.93]) to traditional (AUC=0.93[0.92-0.95]) and hybrid (AUC=0.96[0.94-0.97]) predictive models in training cohorts. However, the deep learning model handled ambiguous tests more cautiously than other models; the deep learning model was more likely to designate an ambiguous test as inconclusive (odds ratio=4.21[2.78-6.38]) versus traditional/hybrid approaches.

**CONCLUSIONS:** By considering complex spatial-temporal information beyond conventional anorectal function metrics, deep learning on 3D-HDAM technology may enable gastroenterologists to reliably identify and manage dyssynergia in broader practice.

## INTRODUCTION

Bowel disorders such as constipation and fecal incontinence lead to over 2 million gastroenterology referrals and 700,000 emergency department visits annually in the United States ^1–4^. Several practice guidelines advocate routine, up-front anorectal function testing in patients failing empiric treatments for these conditions such as laxatives and fiber, to distinguish whether further treatment efforts should broadly focus on gastrointestinal transit or rectoanal coordination ^5,6^. The most common disorder of rectoanal coordination is dyssynergia, which is defined as a “failure of coordinated anal relaxation” during defecation ^7^. Identifying dyssynergia is relevant and clinical important broadly in gastroenterology, because dyssynergia is implicated in up to 40% of cases in typical referral populations and preferentially responds to biofeedback physical therapy^8^.

Contemporary anorectal function tests rely on relatively straightforward measures of the rectoanal pressure gradient or anal canal relaxation (using anorectal manometry) or overall function (measured by expulsion of an inflated balloon inserted into the rectum) to define the “presence” or “absence” of dyssynergia ^9^. Unfortunately, this simplified diagnostic paradigm represents the limitations of available technology and contrasts with the synchronized physiology required to achieve successful defecation that is far more complex than any single binary metric can capture ^10,11^. These problems are recognized in the recent (and first) iteration of the London consensus protocol intended to standardize anorectal function testing: dyssynergia represents only a “minor disorder” due to challenges in stratifying pathologic dyssynergia from physiologic dyssynegia in non-constipated healthy individuals using existing metrics and technology ^7,11,12^. As a result of high variability in the performance and interpretation of anorectal function tests in practice, adoption and acceptance of anorectal function testing in gastroenterology remains poor ^13^.

High-definition anorectal manometry (3D-HDAM) is a recent development providing three-dimensional spatiotemporal mapping of the anorectal canal during simulated defecation in fine detail ^13^. Deep learning is a recent methodologic advancement in artificial intelligence that is capable of discovering important, nonintuitive relationships in complex datasets ^14^. Unlike traditional artificial intelligence approaches that inherently require existing heuristics to guide the resulting algorithms (such as conventional anorectal function metrics in evaluating dyssynergia, adenomatous classification systems for polyps, or dysplastic classification systems for inflammatory bowel diseases or Barrett’s disease) ^15,16^, deep learning harnesses multi-layered artificial neural networks to evaluate non-intuitive patterns and is thus particularly well-suited to complex 3D-HDAM data. As a result, deep learning circumvents the limitations of fitting conventional anorectal function concepts onto 3D-HDAM technology. Deep learning is also capable of learning from real-time video of defecation mechanics provided on 3D-HDAM—recognizing the potential importance of the *act* of defecation during a bowel *movement*. In this study, we aimed to develop and optimize a deep learning algorithm to address the potential unrealized value of 3D-HDAM and facilitate its broader applicability in routinely detecting dyssynergia among patients referred for evaluation of bowel disorders.

## MATERIALS AND METHODS

### Study enrollment

We included consecutive 3D-HDAM procedures performed as part of clinical care at our tertiary care institution between October 1, 2018 and May 13, 2020 across indications (generally constipation and fecal incontinence), recognizing that our intent to develop an algorithm to routinely and automatically interpret defecation maneuvers across clinical indications supersedes a narrower focus on a particular patient group. Our study protocol was approved by our Institutional Review Board.

### Clinical protocol for 3D-HDAM

The 3D-HDAM probe is a pressure sensing catheter utilizes 256 circumferentially placed sensors (16 per level, 16 levels) to record pressures along the length of the anal canal. Procedures were conducted according to a modified London consensus protocol using the ManoScan™ AR 3D HRM system (Medtronic, Minneapolis, MN, USA) in the lateral decubitus position following administration of a tap water or saline enema, recognizing both the limited feasibility of seated manometry using 3D-HDAM equipment and common use of non-physiologic lateral positioning at most centers ^7,13,17^ (**Supplement**).

### Gold standard definition of dyssynergia

Deep learning models algorithmically construct predictors without human intervention, but these approaches still require minimal input to at least define an “abnormal” test. We aimed to define our gold standard for dyssynergia according to London consensus definitions while also accommodating technical nuances of 3D-HDAM technology and avoiding areas of diagnostic controversy in our initial work.^18^ For each simulated defecation maneuver, we defined the technical criteria to classify *no dyssynergia* as ≥20% reduction in the minimum averaged 1-second relaxation within the anal canal over a 15 second defecation period compared to the pre-maneuver averaged 3-second resting pressure. The 20% threshold was used to accommodate increased pressure sensitivity of 3D-HDAM equipment. Studies with paradoxical contraction of the anal canal were classified as *dyssynergia* (corresponding to Rao classification subtypes I-II). Studies with 0-20% relaxation were classified for our study purposes as *ambiguous* (corresponding to Rao classification subtypes III-IV) and reserved for further validation. The intent of reserving these studies was to train the initial algorithm on clearly normal or abnormal studies, and then to evaluate the ability of our algorithm to avoid overfitting dyssynergia classifications when tested in a real-world setting. These definitions are consistent with both the London consensus and Rao classification paradigms ^7,9^.

### Data extraction and preprocessing

Tabular data were imported from ManoView™ software (Medtronic, Minneapolis, MN, USA) into Python-based Jupyter Notebooks. These data from *T* individual timepoints were reshaped into a 16*x*16 matrix (i.e. HDAM image) to form a *Tx*16*x*16 array (i.e. HDAM video) at ten frames per second. To accommodate pressure drift, we subtracted out the averaged 2-second resting pressures for each sensor just prior to each simulated defecation maneuver using metadata accompanying each study.

### Traditional machine learning models

We first developed “traditional” supervised and unsupervised machine learning approaches, which focus on identifying patterns in still images to develop potential predictors of dyssynergia from complex 3D-HDAM data (**Figure 1a-b**). Using these predictors, we trained four supervised and unsupervised machine learning models. The supervised machine learning approaches included logistic regression and linear and quadratic discriminant analysis; parameters were estimated for the manually extracted predictor set (**Figure 1c**). The unsupervised modeling approach involved Gaussian mixture models, a method that determines the probability of having dyssynergia ranging from 0-100%. All of these traditional modeling approaches were trained on the training set and evaluated on the test set. Technical description of these methods is detailed in the **Supplement**.

**Figure 1:**
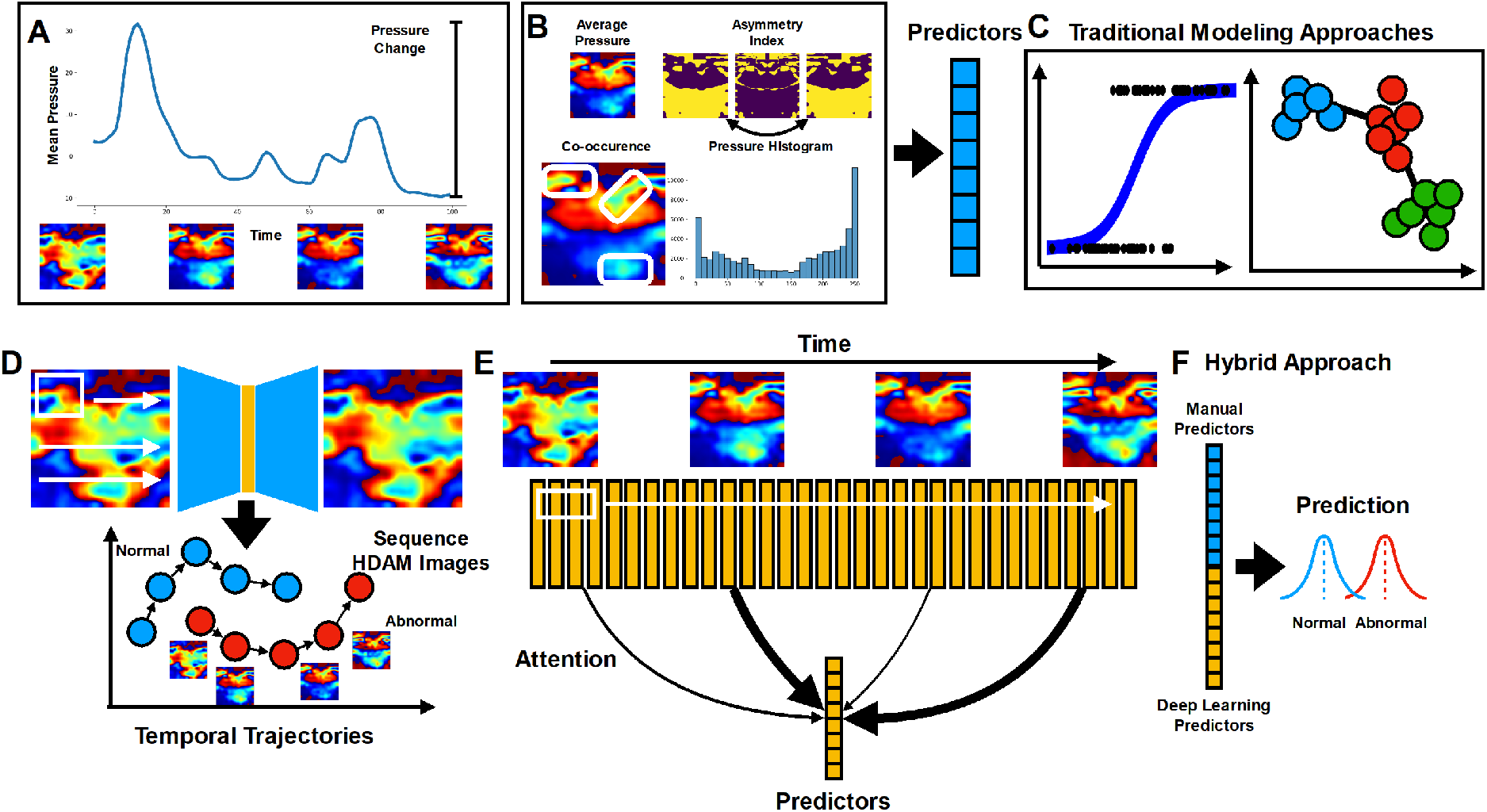
Description of traditional (Panels A-C), deep learning (Panels D-E), and hybrid approaches (Panel F). (A) Example of mean pressure over time a for 10-second simulated defecation maneuver. (B) Predictors are extracted using manually specified heuristics. (C) Predictors are used to fit traditional machine learning approaches. For the deep learning approach, (D) HDAM image frames are compressed using variational auto-encoders (circles) that relate to other images in maneuver akin to a temporal trajectory (arrows) to form patterns. (E) HDAM images in video are integrated using “convolutions” and “attention” to form one set of algorithmically derived predictors. (F) The hybrid approach combines traditional and deep learning acquired predictors.

### Deep learning model

We designed a deep learning approach to evaluate spatial pressure measurements of pressure over time during simulated defecation as a video (**Figure 1d-e**). The entire model was trained end-to-end to make predictions on sequences of HDAM images, model the temporal dependencies between HDAM images, filter out unimportant HDAM images/subsequences, all while simultaneously ensuring spatial features extracted from each HDAM image may be readily incorporated temporally. Model design is further detailed in the **Supplement**.

### Hybrid model

Each modeling approach (machine learning and deep learning) potentially offers a set of complementary predictors that could strengthen the ability to predict abnormal manometry tests. Common predictors between the traditional and deep learning models were first identified using canonical correlation analysis. The hybrid model combined predictors from both the traditional and deep learning models, where predictors outside of the common predictors could potentially provide a boost in predictive performance.

### Assessing diagnostic accuracy of machine learning models

Tests were split at the patient level using grouped 5-fold cross validation splits, ensuring that within-patient squeeze maneuvers were placed in the same training, validation and test sets to ensure that the efficacy of the algorithms were not overstated by mixing statistically dependent samples. Within each cross-validation fold, each study was decomposed into four simulated defecation maneuvers/HDAM videos (i.e. one video for each simulated defecation maneuver). Overall performance was assessed using the Area Under the Receiver Operating Characteristic curve (AUC) on the video-level. AUCs were compared between modeling approaches through a 1,000-sample non-parametric bootstrap sampling of AUCs within held-out sets of cross-validation folds, then calculating pairwise differences in AUC between different algorithms given the same sampling of HDAM data per bootstrap with 95% confidence intervals (CI).

Overfitting is a common problem in machine learning approaches that external validation datasets alone do not fully resolve. To estimate differences in ideal performance compared to real-world performance, we trained our models on “ideal” cases of clear dyssynergia or no dyssynergia and reserved ambiguous tests in a separate “real-world” analysis (i.e. how well each modeling approach tackled ambiguity based on the distribution of abnormal probabilities assigned to held-out ambiguous cases in each test fold). The expectation was that abnormal probabilities for ambiguous cases should coalesce near intermediate probabilities (i.e. 0.5) rather than probabilities that would indicate a more definitive diagnosis (i.e. 0 or 1). This hypothesis was tested through the estimation of odds ratios to assess whether assignment of an ambiguous probability (i.e. closeness to 0.5) actually reflected whether the test was actually ambiguous. These tests were conducted for each modeling approach (traditional, deep learning, hybrid) and compared to see which model had the highest odds ratio, where an odds ratio greater than one indicates the ability to capture and communicate ambiguity.

## RESULTS

### Cohort Characteristics

We assessed 1,208 simulated defecation maneuvers from 302 patients undergoing 3D-HDAM with a mean age of 55.1 years and 243 women (80.5% of the total cohort). 446 maneuvers were reported as ambiguous and reserved for further validation. Excluding ambiguous cases, 59.4% of maneuvers met criteria for dyssynergia (453 of 762 total maneuvers).

### Diagnostic accuracy of deep learning, traditional, and hybrid models to interpret 3D-HDAM to detect dyssynergia

Excluding ambiguous cases, our deep learning, traditional (logistic regression-guided machine learning), and hybrid models achieved comparable performance in detecting dyssynergia (**Table 1, Figure 2, Supplementary Figure 2**) and were reliably able to stratify studies on particular features associated with defecation (**Supplementary Figures 3-5**), visualized using 2D UMAP plot summaries of the features extracted by the deep learning model and for the traditional modeling approaches ^19^. Logistic regression was the most accurate traditional machine learning approach, compared to discriminant analysis and unsupervised approaches (**Table 1, Supplementary Tables 1-2, Supplementary Figure 6**).

**Table 1:**
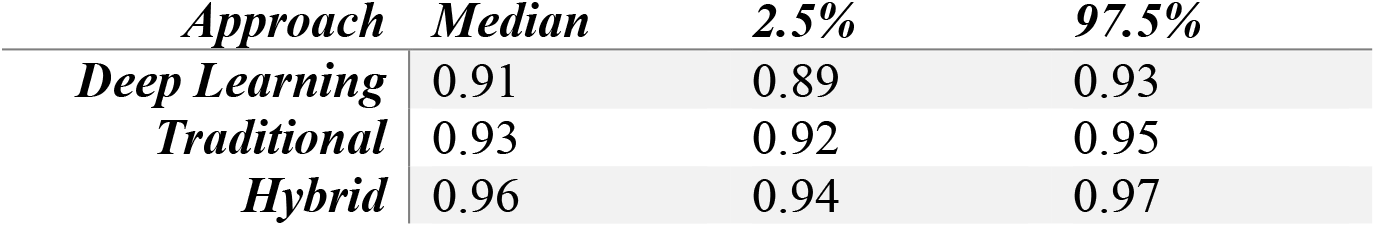
Diagnostic accuracy of deep learning, traditional machine learning, and hybrid models. Results are reported for 5-Fold cross-validated area-under-the-curve (AUC) estimates bootstrapped for each modeling approach. Deep learning, traditional and hybrid machine learning models show an AUC>0.9 in detecting dyssynergia among individual simulated defecation maneuvers.

| *Approach* | *Median* | *2.5%* | *97.5%* |
| --- | --- | --- | --- |
| *Deep Learning* | 0.91 | 0.89 | 0.93 |
| *Traditional* | 0.93 | 0.92 | 0.95 |
| *Hybrid* | 0.96 | 0.94 | 0.97 |

**Figure 2:**
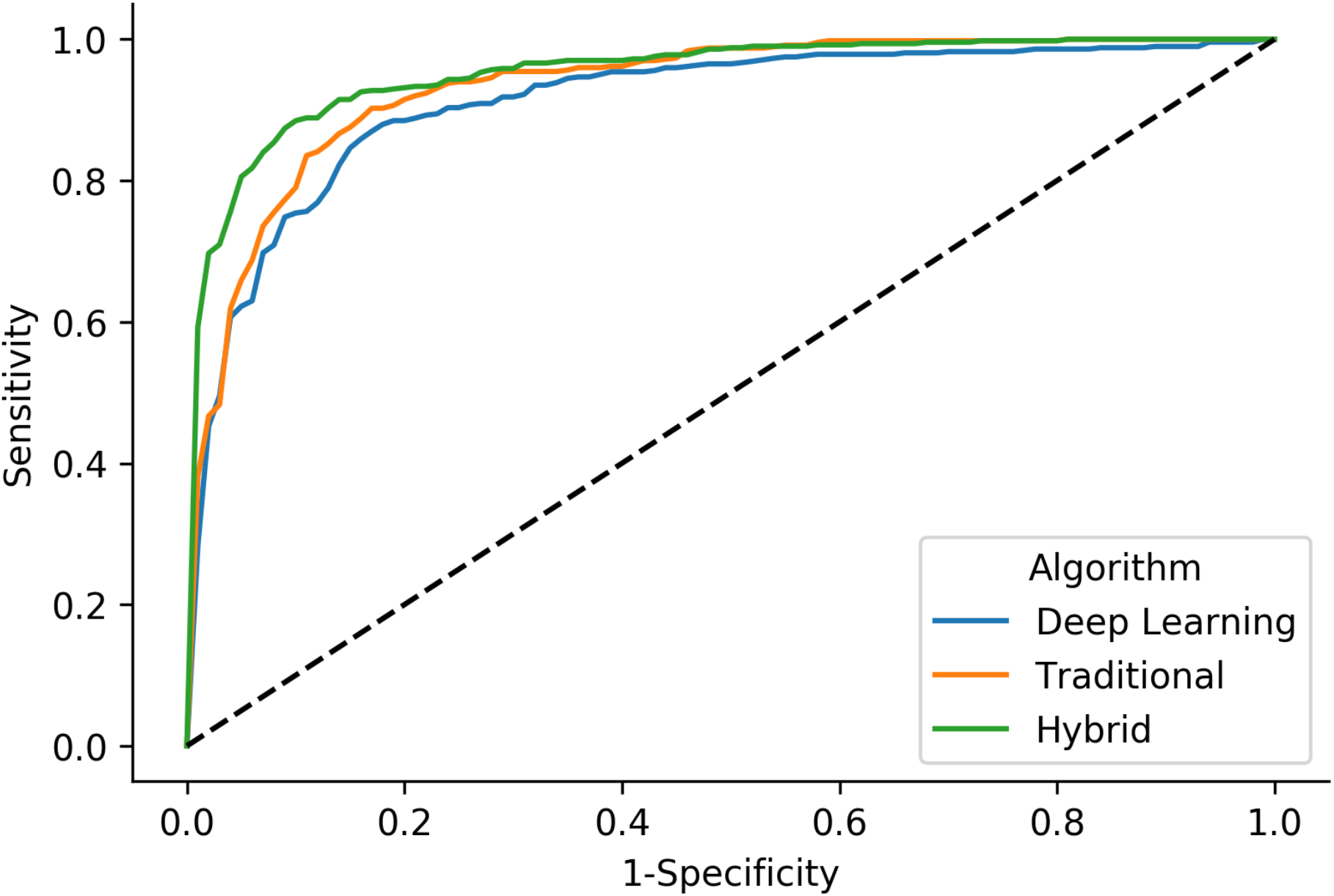
**Receiver-operating characteristic curve (ROC) on the diagnostic accuracy of deep learning, traditional machine learning, and hybrid approaches to detecting dyssynergia**.

Incorporating the ambiguous tests, the deep learning model was more likely to assign an intermediate probability to an ambiguous test (**Supplementary Table 3**), given the conservative nature of how the model estimated the presence of dyssynergia, compared to traditional and hybrid models which tended to more readily stratify maneuvers into dyssynergia (i.e. overfitting). This suggests the ability of the deep learning algorithm to effectively communicate the ambiguity of the case despite not having been exposed to prior examples. These results were further corroborated in density plots of the abnormal probabilities (**Supplementary Figure 6; Supplementary Table 4**) for the ambiguous cases and in embedding plots (**Supplementary Figure 4**), where ambiguous cases were more likely to be near the decision boundary for the deep learning model (**Supplementary Table 3**).

### Exploring determinants of dyssynergia in deep learning and traditional models

Inherent to rationale for using deep learning methodology, these methods are capable of identifying complex determinants of dyssynergia but are not capable of fully describing the nature of these determinants. Addressing the opaqueness of how these algorithms actually function is increasingly important to physicians, patients, and regulators ^20^. Identifying correlations between determinants of dyssynergia in deep learning compared to traditional model predictors serves two purposes: (1) to exclude factors which are unlikely to explain differences in performance and (2) to validate that the deep learning approach at least considers some conventional concepts associated with dyssynergia to test content and construct validity. With manual interpretation, dyssynergia is defined by the rectanal pressure gradient and anal relaxation based on changes in average or maximum pressures aggregated across all sensors along an anorectal catheter ^7^. As described in our methods, these factors were incorporated directly into development of our traditional machine learning models. Our canonical correlation analysis shows that these factors are important to several—but not all— deep learning predictors, suggesting other facets of deep learning that are able to meaningfully stratify studies beyond conventional anorectal function metrics (**Figure 3**). On one hand, the overlap across several determinants in both models emphasizes that deep learning and the traditional machine learning techniques indeed acquire similar predictors representative of dyssynergia (**Supplementary Figure 8**). Further analyses using SHAP plots identified differences in the ranked importance of specific determinants in traditional and hybrid models, suggesting that deep learning approaches (embedded in the hybrid model) offer additional nuances not found by manual predictor extraction (**Supplementary Figure 9**).

**Figure 3:**
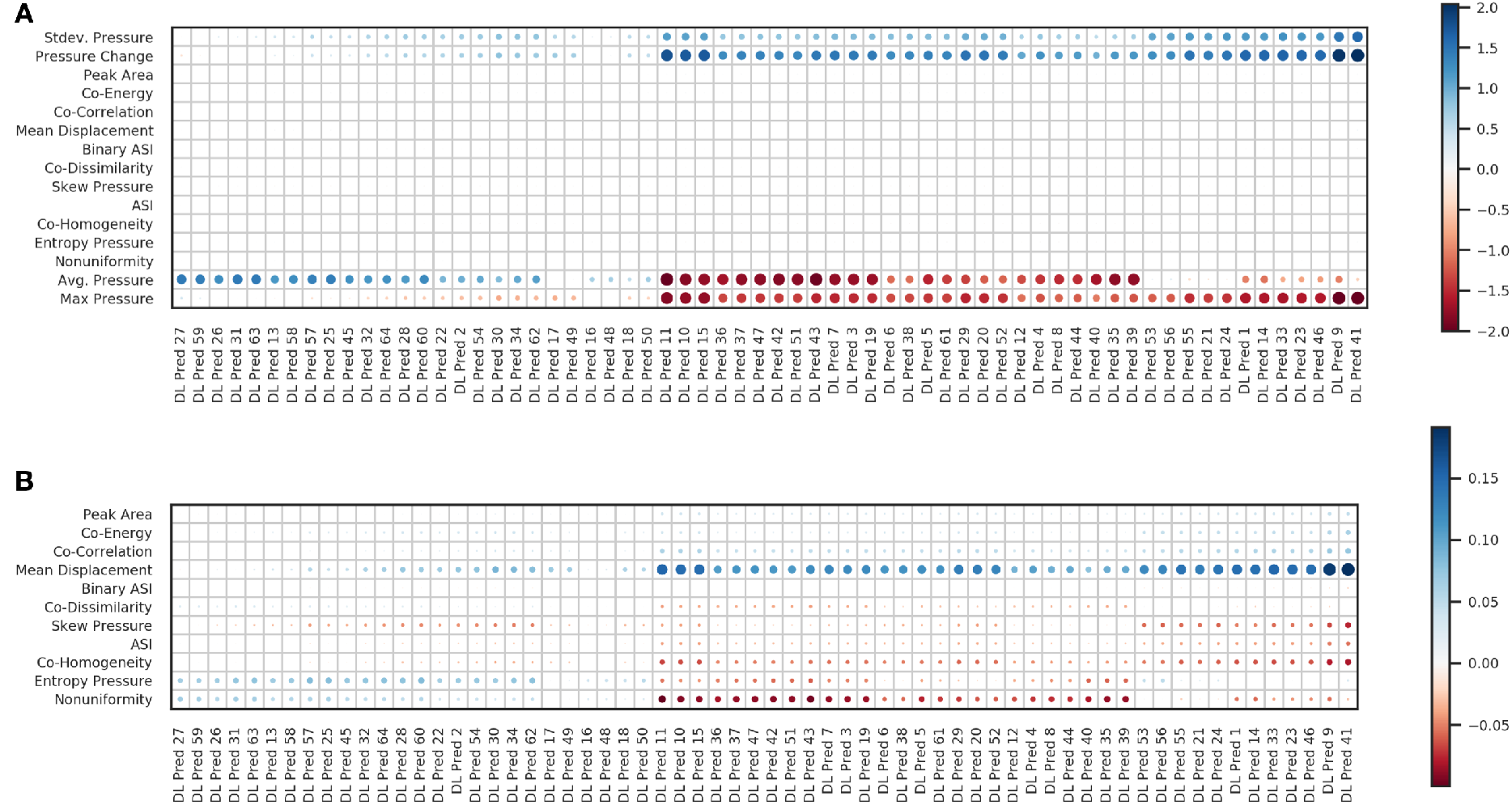
Canonical correlation analysis heatmap relating determinants of dyssynergia by comparing derived test-level deep learning predictors to traditional machine learning approaches. By nature of deep learning methodology, the important, non-intuitive features themselves are unknown and only distinguishable using the gradient and attention methods that highlight HDAM information on a test-by-test basis. However, colors trending toward blue (positive) and red (negative) on the figure signify overlapping associations with at least some of the traditional machine learning determinants of dyssynergia. The magnitude of each association is proportional to the size of the circles. Predictors prepended with “Co-” were generated from the gray-scale co-occurrence matrix. (A) Relationships for all traditional predictors versus the deep learning predictors. Associations for four pressure-based traditional predictors are of far greater magnitude than the other predictors, so these four predictors were removed in (B) to reveal more nuanced relationships. Note how deep learning derived predictors (denoted *DL Pred*) 16-18 and 48-50 do not appear to have relationships with other traditional predictors, yet deep learning predictor 48 is the most important predictor of the hybrid model, determined via the SHAP plots in the supplementary material.

### Identifying 3D-HDAM images representative of dyssynergia using deep learning

While the specific factors driving deep learning are unknown, attention weights can be calculated to at least demonstrate representative HDAM images which are important to the deep learning algorithm in stratifying dyssynergia from no dyssynergia (**Figure 4**). We also report representative real-time 3D-HDAM videos to characterize regions of the anal canal during manometry compared to composite pressure readings which are common in current practice, based on the presence or absence of dyssynergia classified using deep learning.

**Figure 4:**
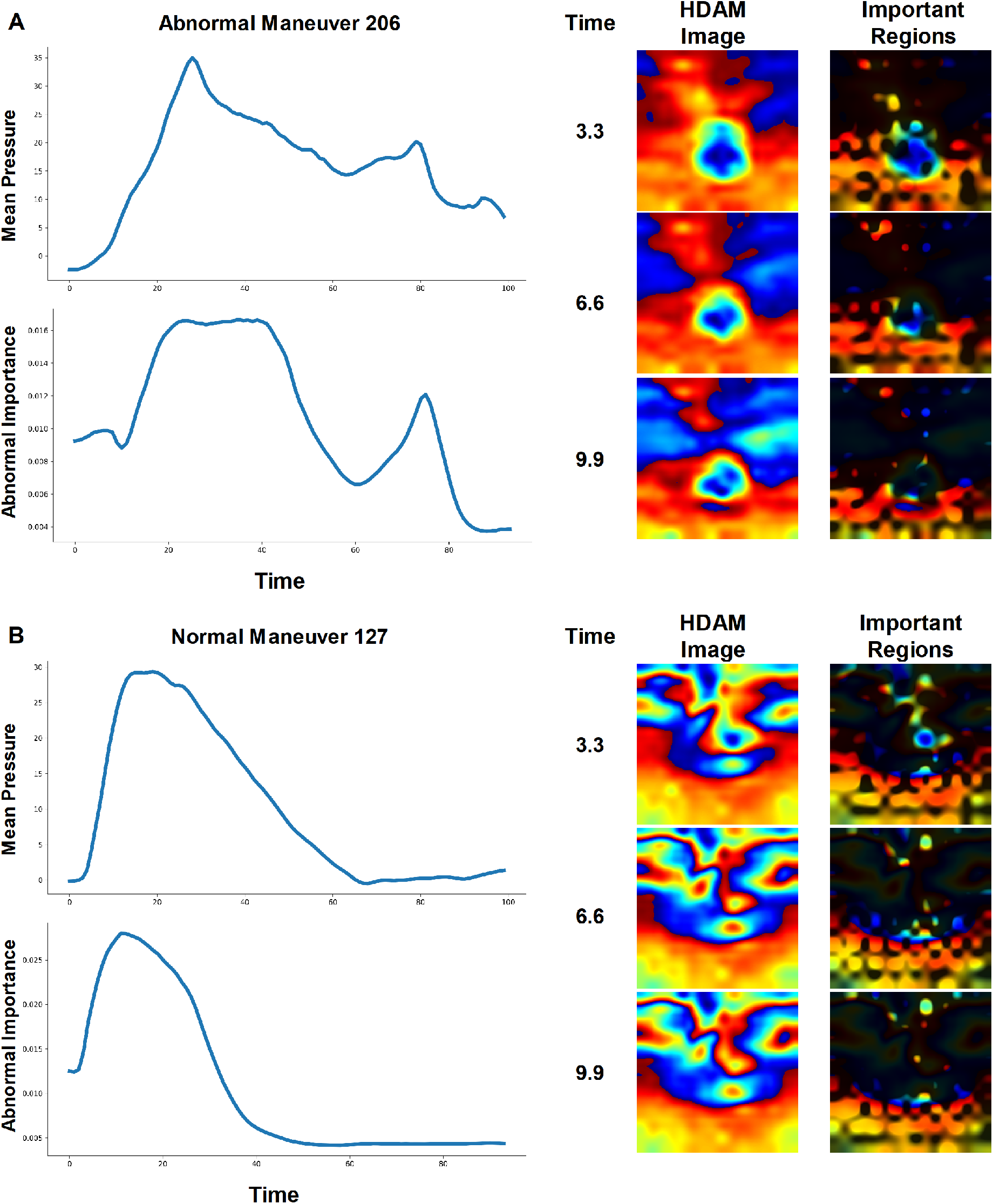
Representative 3D-HDAM images identify spatial and temporal features characteristic of: (A) dyssynergia or (B) no dyssynergia, as classified by deep learning algorithms at three representative time-points during each 10-second simulated defecation maneuver. Averaged composite pressures during simulated defecation (upper-left graphs on both panels) represent typical pull-through pressure plots during conventional anorectal manometry. Attention scores (lower-left graph of both panels) are reported in milliseconds for each 3D-HDAM image in a 10-second real-time video during simulated defecation; higher attention scores suggest subsequences important for detecting dyssynergia. On right side of panels, spatial features have been retained in images particular timepoints to indicate their importance for prediction. Animations of important spatial-temporal features corresponding to these tests have been provided in the supplementary videos.

## DISCUSSION

We developed a novel deep learning algorithm to interpret simulated defecation maneuvers using 3D-HDAM technology to facilitate a deeper evaluation of defecation mechanics in a manner that appears capable of improving the generalizability of 3D-HDAM in broader gastroenterology practice. We show that deep learning can uncover important, nonintuitive patterns in complex defecation mechanics and distill those findings to better integrate the multiple parameters from ARM that do not yet have clinical utility (i.e. spatial and temporal patterns from raw input image data which do not correlate with traditional HDAM features). We also show that deep learning appears superior to previous machine learning models and may be appropriate to overcome the limitations of applying conventional anorectal function metrics to the complexities of both 3D-HDAM technology and anorectal physiology. Importantly, our algorithm can be adapted as standardized protocols to interpret anorectal manometry evolve.

Bowel disorders including constipation and fecal incontinence are a frequent reason for referral to gastroenterologists ^1^. Anorectal function testing is recommended as the next best step for patients failing empiric treatment (such as laxatives or fiber) ^5,6^. Yet only an estimated 2% of patients meeting Rome IV criteria for chronic constipation have ever undergone anorectal function testing ^21^. For the relatively few patients who undergo anorectal function testing in broader practice, a recent international survey of the relatively few available centers offering anorectal function testing found significant variation in testing protocols and interpretation methods ^13^. To accommodate these problems, the London consensus protocol was recently developed to standardize and define best practices for anorectal function testing ^7^. Despite this accomplishment, no single test or metric has been deemed sufficient to diagnose dyssynergia. Furthermore, dyssynergia is frequently found in healthy populations using conventional anorectal metrics ^22^. While concern for dyssynergia is one of the most frequent reasons to refer adult patients for anorectal function testing, dyssynergia in itself remains a “minor disorder” in the London consensus protocol as a result of these diagnostic dilemmas.

A likely explanation for these gaps between knowledge and practice is the stark contrast between the simplicity of available metrics used to evaluate defecation compared to the well-known complexity of defecation mechanics. 3D-HDAM provides a detailed spatio-temporal evaluation of pressure along the anorectal canal and offers the promise of overcoming structural and functional complexities of defecation. However, 3D-HDAM technology only translates a complex medical problem into a complex image, and physician interpretation is still required (similar to cross-sectional imaging or endoscopy). Without appropriate standards for interpretation, variation is inevitable. As in other areas of gastroenterology in which an accurate diagnosis essentially relies on image processing, we have shown that deep learning methods are capable of distilling complex anorectal physiology to fit the needs of the clinician in determining whether to direct a patient to biofeedback physical therapy.

Deep learning represents a class of computational heuristics that represent images (or in this case pressure sensor readings that may be spatially correlated) as a set of *nodes*, from which non-obvious features extracted from an image can be integrated to form patterns ^14^. What sets deep learning approaches apart from other heuristics is its ability to construct patterns that the model itself deems to be most relevant for prediction, in contrast to approaches that rely on conventional anorectal metrics. As such, deep learning approaches may potentially offer additional insight on factors important to manometry assessment that conventional anorectal metrics cannot. Furthermore, deep learning adds the capability of evaluating real-time videos of simulated defecation rather than still images alone, recognizing the differences in clinical needs between evaluating simulated defecation using deep learning and identifying adenomatous polyps or dysplastic lesions (that do not evolve over months-to-years rather than over the course of endoscopy) for which still-image processing using traditional machine learning methods may be sufficient ^15,16^.

There are several limitations which we considered in our study design, recognizing that our intent was to evaluate the potential generalizable clinical impact of an appropriate deep learning approach and not to resolve contentious nuances in the tertiary care evaluation of anorectal disorders. Our study includes a cohort of individuals referred for constipation and/or fecal incontinence, but does not include asymptomatic individuals. There is a lack of consideration for the results of balloon expulsion testing in our present study, recognizing that balloon expulsion may be the more relevant test in current practice (a finding that is outside the scope of our present study question).^12,18^ We did not utilize rectal balloon pressure in our proof-of-concept algorithm, largely due to variation in rectal balloon pressures using 3D-HDAM equipment and technical design differences between the rectal balloon on 3D-HDAM catheters compared to other manometry catheters ^23^. To address this, we note that rectal pressures are inherently included in the proximal portion of the anal probe and also that these pressures are not presently required by the London consensus protocol. Further efforts to refine the model in a nationwide multi-center collaborative are ongoing, recognizing (1) the novelty of developing artificial intelligence applications for motility disorders and rapidity of ongoing advancements in deep learning methods, (2) our focus on evaluating the performance of deep learning on real-world ambiguous studies in a separate validation set, and (3) our use of machine learning/hybrid approaches and SHAP plots to shed light on the inherent “black box” nature of our deep learning algorithms.

In summary, we developed a deep learning algorithm capable of interpreting 3D-HDAM to assess for dyssynergia. Our algorithm can be adapted as future iterations of the London classification system evolve. These novel findings offer the broad potential for gastroenterologists to have access to relatively limited neuro-gastroenterology testing modalities such as anorectal manometry—deep learning algorithms on 3D-HDAM technology offers the potential to resolve limitations of current technologies and paradigms to more effectively distill complex physiologic anorectal function assessments into simple, accessible clinical recommendations on appropriate care. Our findings provide hope for the 98% of individuals with chronic constipation and other bowel disorders who have never undergone anorectal function testing and advocate the need for further efforts to encourage a proper evaluation of anorectal function given the prevalence and burden of these disorders in practice.

