## Supplementary Material for "Video-Based Deep Learning to Detect Dyssynergic Defecation with 3D High-Definition Anorectal Manometry"

### EXPLANATION OF TRADITIONAL SUPERVISED AND UNSUPERVISED MACHINE LEARNING PREDICTORS AND APPROACHES

The following common predictors were built off of HDAM videos defining *traditional* features:

- **Tabulated statistics derived from pressure histogram of HDAM image after averaging across all time points, including:** 1) average pressure level, 2) standard deviation or variation of pressure levels, 3) smoothness, a measure to detect the sharpness of transition from low to high pressure in histogram, 4) third moment, indicating skewness of histogram, 5) uniformity, whether pressure across channels was similar, and 6) entropy, measurement of disorder in pressure values.
- **Symmetry:** Assymetric index (ASI) calculated by subtracting differences between sensors corresponding to a folded HDAM image across a vertical median line.
- **Pressure Changes:** Minimum pressure for the HDAM video subtracted from maximal pressure, significant changes in pressure could signify abnormalities.
- **Grey-Level Co-occurrence:** Here, the HDAM image taken at maximum pressure during the maneuver is converted into a co-occurrence matrix that examines how frequently pairs of surrounding pixels occur in other parts of the image (pairs being defined by the current and adjacent pixel by a diagonal, horizontal or vertical position). From this co-occurrence matrix, metrics are calculated based on: 1) the extent to which pixel pairs correlate with one another, 2) energy (i.e. the sum of the square elements), and 3) homogeneity (i.e. the extent of similarity among off-diagonal elements of the co-occurrence matrix compared to on-diagonal elements). Additional texture features were calculated as previously reported in prior informatics approaches.

To serve as a baseline comparison to the machine learning approaches, we generated predictions using the following supervised machine learning algorithms:

- **Logistic Regression:** Part of the family of generalized linear models and widely used across medicine and epidemiology, this modeling approach assumes that the outcome $Y$ follows a binomial distribution with probability $p$, conditional on the predictors, $\vec{X}$, via the relationship $E[Y|X] = P(Y = 1|\vec{X}) = p = \frac{1}{1+e^{-\vec{\beta}\cdot\vec{X}}}$.
- **Linear and Quadratic Discriminant Analysis:** Discriminant analyses model a categorical outcome (presence or absence of dyssynergia) according to continuous or interval-type predictors, with the objective that outcomes can be maximally distinguished using linear combinations of the independent variables. This is accomplished by estimating a projection matrix (mathematical linear combination of predictors) that maximizes between-class variance while minimizing within-class variance via eigen decomposition of such covariance matrices. For Linear Discriminant Analysis (LDA), it is assumed that the within-cluster covariance is shared between the classes, while for Quadratic Discriminant Analyses (QDA), this assumption is violated, resulting in a quadratic decision curve. Both LDA and QDA derive their objective function and make predictions based on a likelihood ratio test, which assumes that data from each class are multivariate and normally distributed, thus the likelihood of the predicted class should be larger than that of the other classes by some threshold.

Additionally, we estimated the predictive performance of one select unsupervised clustering technique– Gaussian Mixture Model, which assigns probabilities of cluster membership (which may correspond to normal/abnormal exams). The assigned clusters were optimally aligned to the outcomes of interest in order to compare the predicted probabilities to the presence of an abnormal test. KMeans and hierarchical clustering, other two popular clustering techniques, were excluded from the analysis since they did not offer straightforward mechanisms of assigning cluster assignment probabilities on held-out test data.

**Gaussian Mixture Models:** Generalized version of k-means clustering that assumes data instances arise from a mixture of multivariate gaussian distributions with unknown mean and covariances. Typically fit using variational inference or expectation maximization, which iteratively: 1) estimates the posterior probability of the data instance being generated by the cluster, and 2) fitting the parameters of the multivariate gaussian distributions to the data.

### EXPLANATION OF DEEP LEARNING METHODOLOGY

Artificial neural networks (ANN) represent these complex data as nodes, passing this signal through a series of intermediate layers which calculate complex interactions and non-linear transformation of the predictors, to form an output that is compared to the expected output. To simultaneously capture and integrate spatial-temporal correlations in a manner which represents the probability of an abnormal test, we trained convolutional neural networks (CNN) on individual HDAM images. CNNs slide learnable filters across these pressure sensors to detect patterns and apply pooling operations to integrate spatially dependent patterns to form an abstraction of the image. We utilized a variant of these types of networks called convolutional Variational Auto-Encoders (VAE), which compress HDAM images $\{\overleftrightarrow{X_t}\}_{t\in[1,100]}$ (*Tx16x16* sequence of images) into latent vectors $\{\vec{z}_t\}_{t\in[1,100]}$ (via an encoder neural network) that encapsulate key information while revealing nuances. These "embeddings" are decompressed into their original form $\{\hat{X}_t\}_{t\in[1,100]}$ (via a decoder neural network) to ensure that the latent information contained enough information to successfully recapitulate the original data. The optimization procedure balances the expressiveness of the latent vectors (by assuming a multivariate normal distribution, $\vec{z} \sim N\left(\vec{\mu}, \overrightarrow{\sigma^2}\right)$) with the ability to reconstruct the data. To bias the network towards acquiring features that pertain to abnormal stage, we additionally fed the latent information into additional neural network layer that predicts whether the HDAM image came from an abnormal maneuver, $\hat{y} = f(\overrightarrow{z_t})$, where $P(abnormal|X) = \sigma\big(f(\vec{z}_t)\big)$.

Predicting on the latent vectors of the HDAM image assumes that the images are temporally independent. However, videos contain collections of dependent images, representing a temporal trajectory that must be adequately modeled. Additionally, many of the HDAM images contained in an HDAM video are irrelevant for the prediction, so it would be useful to have a filtering mechanism to remove these irrelevant images. With these design principles in mind, we applied a series of temporal convolutions to share information between adjacent images in the form of a parameterized averaging of the latent information in small windows to capture temporal dependencies between images. Then, irrelevant HDAM images/subsequences were filtered out through an attention mechanism, which learns to assign an importance score to small subintervals in the maneuver based on the algorithm's perception of the subsequence's importance for prediction of an abnormal test. Application of the attention mechanism results in one predictor vector that describes the entire maneuver, $\vec{z} = \sum_{t\in[1,100]} \alpha_t \vec{z_t}$. Finally, this aggregate information is passed to a final classification layer that detects whether the entire maneuver was abnormal, $\hat{y} = f(\vec{z})$, where $P\left(abnormal \middle| \{\overleftrightarrow{X_t}\}_{t\in[1,100]}\right) = \sigma\left(f(\vec{z})\right)$.

For extraction of spatial deep learning predictors, encoder of the VAE is comprised of four convolutional layers (with a stride of two/three to downsample), a reparameterization layer (mean and standard deviation to "guide" the posterior distribution), followed by a series of four upsampling convolutional layers, each followed by convolutions. Batch normalization and a leaky ReLU activation function was applied after each convolution layer. The auxiliary classifier for regularizing the VAE was a linear layer that projected 512 nodes of the posterior down to two dimensions.

The architecture of the convolutional variational autoencoder is as follows, motivated by coarse architecture search for architectures that yielded lowest validation loss and inspired by design put forth by MICCAI educational initiative:

1. Encoder:
    a. Input layer, image is of 1x16x16 dimensions.
    b. Application of convolutional layer (2D-Convolution, Batch Normalization and Leaky ReLU activation) with 32 1x3x3 filters, a

stride length of 2 and padding of 1. Resulting feature maps are of dimensionality 32x9x9.

c. Convolutional layer, 128 32x3x3 filters, stride of 1, padding of 1, with resultant image shape of 128x9x9.
d. Convolutional layer, 512 128x3x3 filters, stride of 1, padding of 1, with resultant image shape of 512x4x4.
e. Convolutional layer (no Batch normalization), 512 512x4x4 filters, stride of 1, padding of 1, with resultant image shape of 512x1x1.
f. Two fully connected hidden layers of with input and output size of 512.

2. Reparameterization layer: Two fully connected hidden layers of with input and output size of 512. First layer represents
3. Decoder:
    a. Transposed convolution layer (similar to convolution layer but with upsampling rather than downsampling), maps 512-dimensional vector to 128-dimensions and upsamples to 128x4x4 image.
    b. Convolution retain 128x4x4 image.
    c. Transposed convolution layer, upsamples to 64x8x8 image.
    d. Convolution maps from 64 to 32 channels, resulting 32x8x8 image.
    e. Transposed convolution upsamples to 32x16x16 image.
    f. Final convolution maps from 32 to one channel, returns 1x16x16 image.
4. Auxiliary Classifier: Fully connected layer from encoded 512 dimensions to 2 dimensions (HDAM image level) representing probability for abnormal.

For extraction of temporal features, 512-dimensional vectors were extracted from the VAE, correspondent to each time point, forming for each HDAM video, a $Tx512$ dimensional matrix. The temporal convolutional layers integrated information from a neighborhood of 7 timepoints and mapped the 512-dimensional vectors to 32-dimensional vectors (given the kernel size of 7, resulting matrices were $(T-6)x32$ dimensional). The attention layers transform the 32-dimensional deep learning features at each timepoint into a gating score (via linear layers and sigmoid transforms) that serves to prune information from irrelevant timepoints. Dyssynergia-specific attention scores undergo a Softmax transformation to normalize the importance scores, which are then multiplied by the 32-dimensional embeddings to prune the irrelevant data, yielding one 32-dimensional predictor which encapsulates the HDAM video. A second set of 32-dimensional predictors were generated, which were based on attention scores specific to normal tests, but were not the focus of this work. A total of 64 deep learning predictors (the activation of specific nodes are denoted by *DL Pred* in the manuscript and supplemental figures) were estimated, which utilized the two sets of 32-dimensional vectors were partitioned, each serving as input for normal and abnormal classification layers.

Finally, final classification layers transform the two 32-dimensional embeddings into a 2-dimensional vector which represents the logits, which may be further transformed via a logit/Softmax link function to yield the Dyssynergia-specific probabilities.

Model parameters were updated via gradient descent with adaptive moment estimation optimization (Adam) using the PyTorch framework. HDAM images were normalized by mean and standard deviation of the pressure channels prior to input into the model.

The model was regularized using a combination of loss functions. For initializing the HDAM image representation, a combination of reconstruction loss (mean squared error between reconstructed image and original image), KL divergence between sampling the latent predictor space with a known guide posterior distribution, and a supervised objective (cross entropy) on whether the image came from an abnormal maneuver were utilized. The losses were weighted in order to ensure that the magnitude of one of the losses was not given greater emphasis versus the rest of the joint losses. For aggregation to an HDAM video level prediction, only the supervised objective was considered. Supervised objectives gave higher weight to the minority predicted class in order to calibrate the final predictive probabilities from 0-1 rather than establish an ideal cutoff for an abnormal case via a sensitivity analysis. High learning rates were modulated with cosine annealing functions in order to optimize the tradeoff between exploration and exploitation of the loss landscape, ensuring convergence of the validation loss to a series of local minima, and random pruning of HDAM images from HDAM videos during training further improved generalizability of the model. The epoch of the minimum validation loss during training was used as early stopping criterion to select the model for a given training run under the premise that the selected model would generalize to the held-out test dataset. Models were selected based on minimum loss across the validation set of cases before being evaluated on the test set of each cross-validation fold.

**3D-HDAM Protocol**

The handheld 3D-HDAM probe was carefully positioned to capture the entirety of the anal canal while also capturing intrarectal pressure (proximal to the anus) and environmental pressure (distal to the anus); procedures in which the entirety of the anal canal was not assessed were excluded from this study. Relevant to this study on simulated defecation maneuvers, our 3D-HDAM clinical protocol involves four 10-second simulated defecation maneuvers interspersed by 30-second recovery intervals. The fourth simulated defecation maneuver is also routinely performed with rectal balloon insufflation to 30mL of air at our institution and was included in this analysis, recognizing that the diagnostic criteria for dyssynergia are independent of the technical intricacies of maneuver.

Rectal balloon pressures are measured and reported at our institution. For the purposes of this study, we reserved rectal balloon pressures for future investigations, given that the added value of 3D-HDAM primarily relates primarily to the anal canal measurements and to avoid disagreement among experts on the value of rectal balloon pressures outside the scope of this study.

### INTERPRETATION TECHNIQUES

For our traditional and hybrid modelling approaches, we applied the SHAP (Shapley Additive Feature Explanations) interpretability framework to identify important determinants of dyssynergia(19,20). This model identifies predictors using a basic linear model to approximate our complex machine learning algorithm, in which coefficients of this model represent the importance of each predictor. Important predictors were identified by summing the "predictor importances" across all simulated defecation maneuvers in our cohort (**Supplementary Figure 1a**).

Unfortunately, traditional machine learning models do little to highlight specific spatio-temporal patterns which are important in considering defecation as a literal bowel movement (video) rather than a still image. We interpreted the full context of the 10s (time) x16x16 HDAM video array in detecting dyssynergia using two methods: gradient-based backpropagation methods and attention. The gradient-based backpropagation methods identify important spatial patterns in an HDAM image by first perturbing the output and then propagating this information backwards through the neural network layers. This approach may be integrated with visualization of the attention scores, which temporally illustrate important HDAM maneuver subsequences when visualized (**Supplementary Figure 1b-c**).

**Supplementary Figure 1: Illustration of interpretation techniques to identify potentially important predictors and patterns in our machine learning models.** (A) Shapley Additive Feature Explanations (SHAP) was applied to identify important predictors in our traditional model. Our deep learning model was also interpreted using (B) guided backpropagation and (C) *Attention Over Time* to reveal potentially important HDAM images to predict dyssynergia.

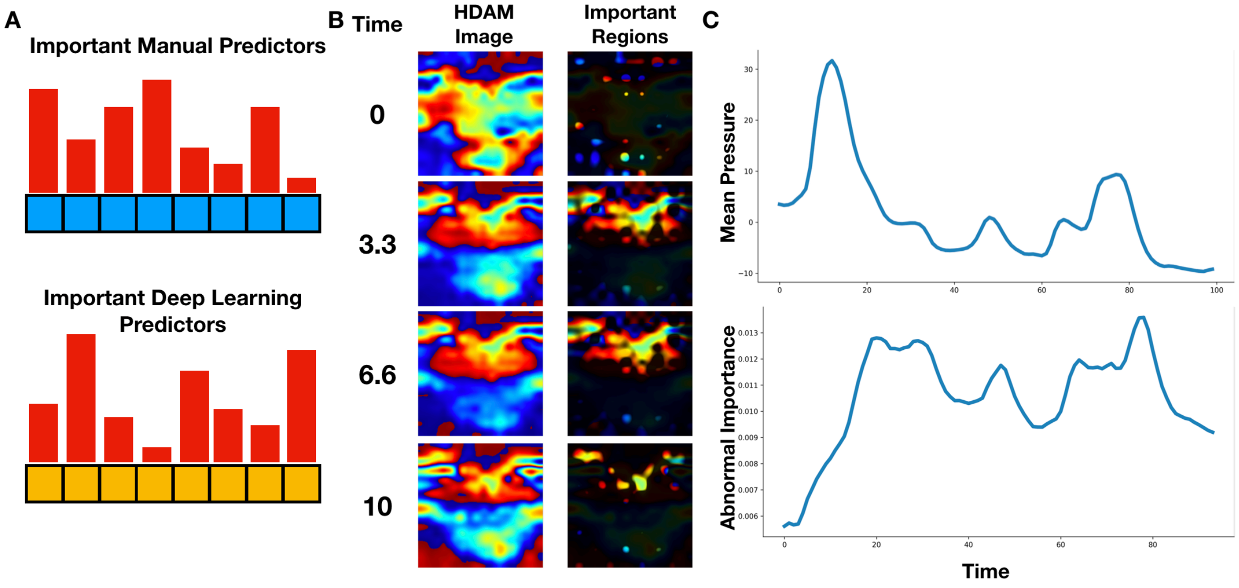

### PERFORMANCE OF MODELING APPROACHES AND LOW DIMENSIONAL EMBEDDINGS

**Supplementary Figure 2: Boxenplots of bootstrapped AUCs averaged across validation folds for deep learning, traditional and hybrid approaches.**

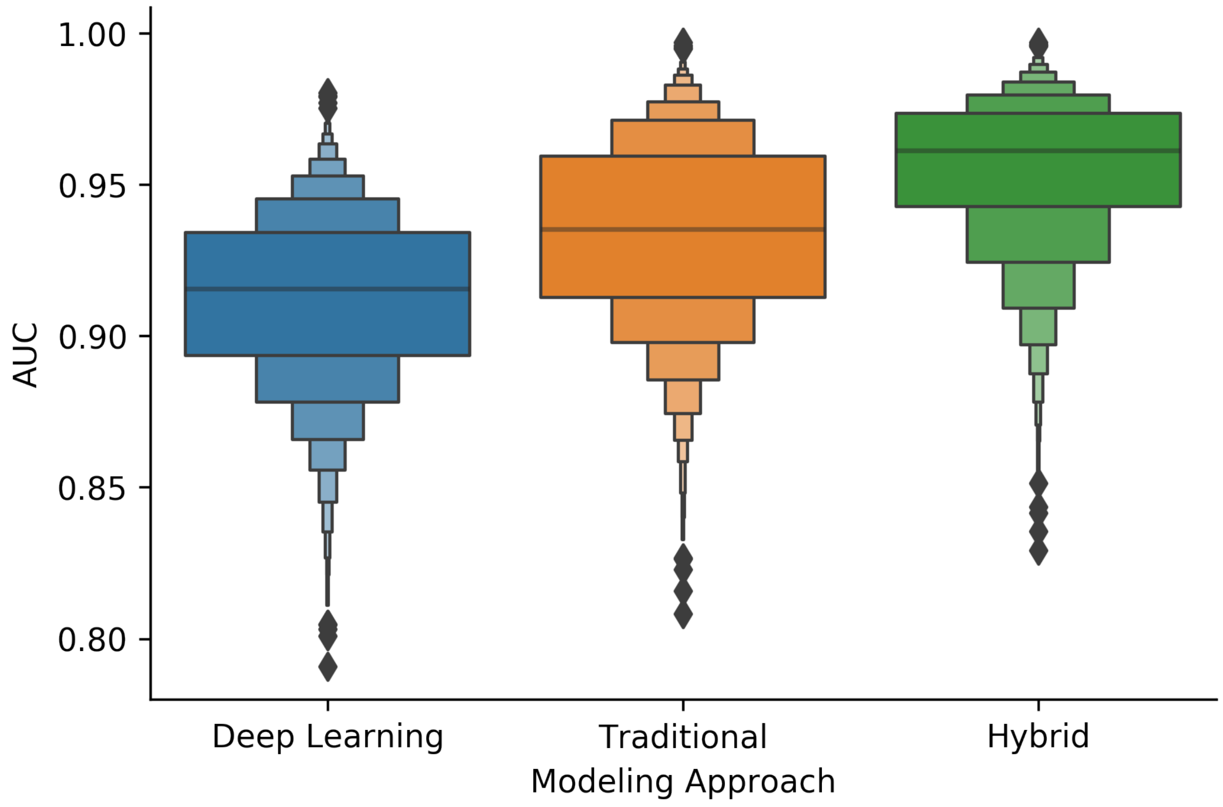


**Supplementary Table 1: Bootstrapped 95% confidence intervals for paired differences in AUCs between modeling approaches.** The AUC of approach 2 is subtracted from the AUC of approach 1 during the bootstrap.

| Approach 1 | Approach 2 | Median Difference | 2.5% | 97.5% |
|---|---|---|---|---|
| **Traditional** | **Deep Learning** | 0.02 | -0.04 | 0.09 |
| **Hybrid** | **Traditional** | 0.02 | -0.02 | 0.07 |

**Supplementary Figure 3: UMAP Embedding plots** for (A) deep learning models, (B) manual feature extraction, and (C) hybrid approach on maneuver-level; normal tests indicated in blue, while abnormal tests indicated in orange; UMAP plots represent high-dimensional data (anything with dimensionality larger than imaginable) in a 2D cartesian coordinate system; this dimensionality reduction technique preserves the relationships between HDAM videos with similar attributes such that videos that may cluster together based on a multitude of traditional and deep learning acquired attributes may cluster similarly in the lower-dimensional space and may be visualized as such

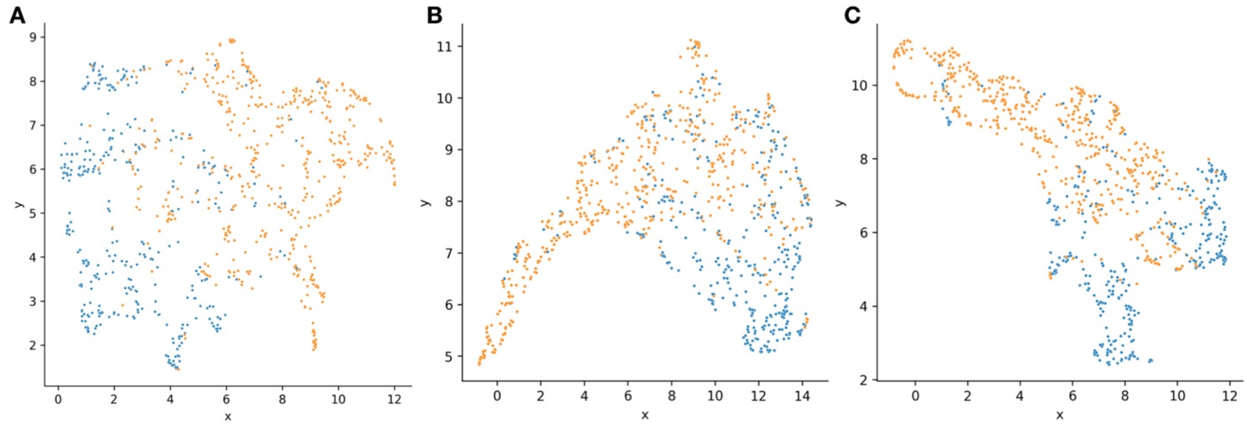


**Supplementary Figure 4: UMAP Embedding plots** for (A) deep learning models, (B) manual feature extraction, and (C) hybrid approach on maneuver-level; normal tests indicated in blue, abnormal tests indicated in orange, and including ambiguous cases, which are colored green

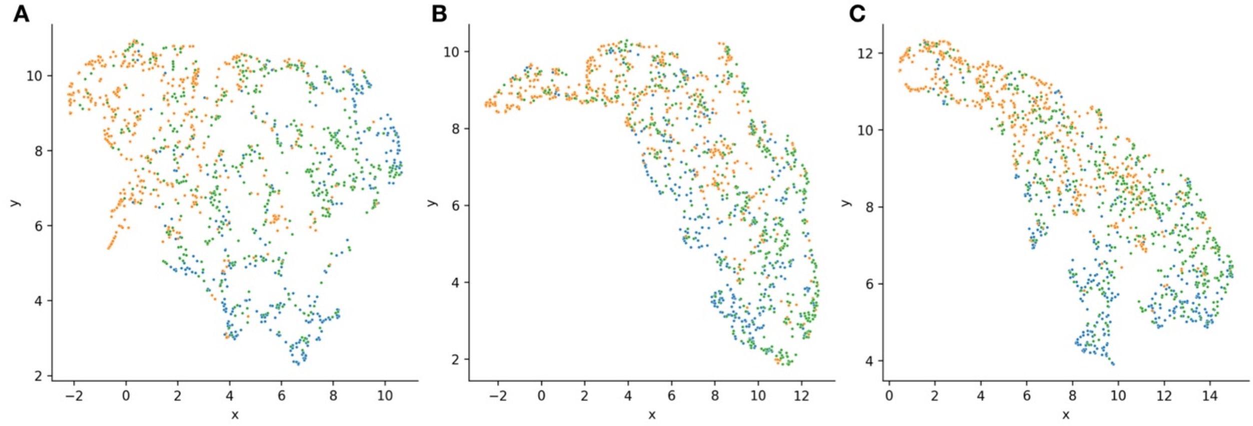

**Supplementary Figure 5**: Embedding plot for deep learning models on HDAM-image level of training set

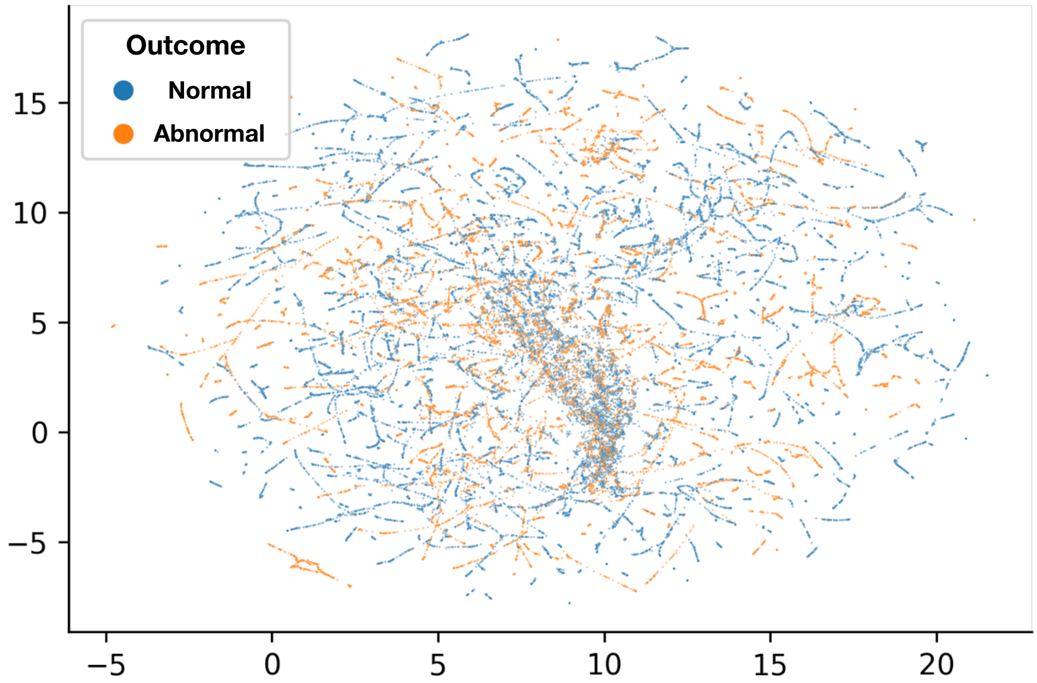

**Supplementary Figure 6:** Boxenplots of bootstrapped AUCs averaged across validation folds for Gaussian Mixture Model, Quadratic Discriminant Analysis, and Linear Discriminant Analysis methods

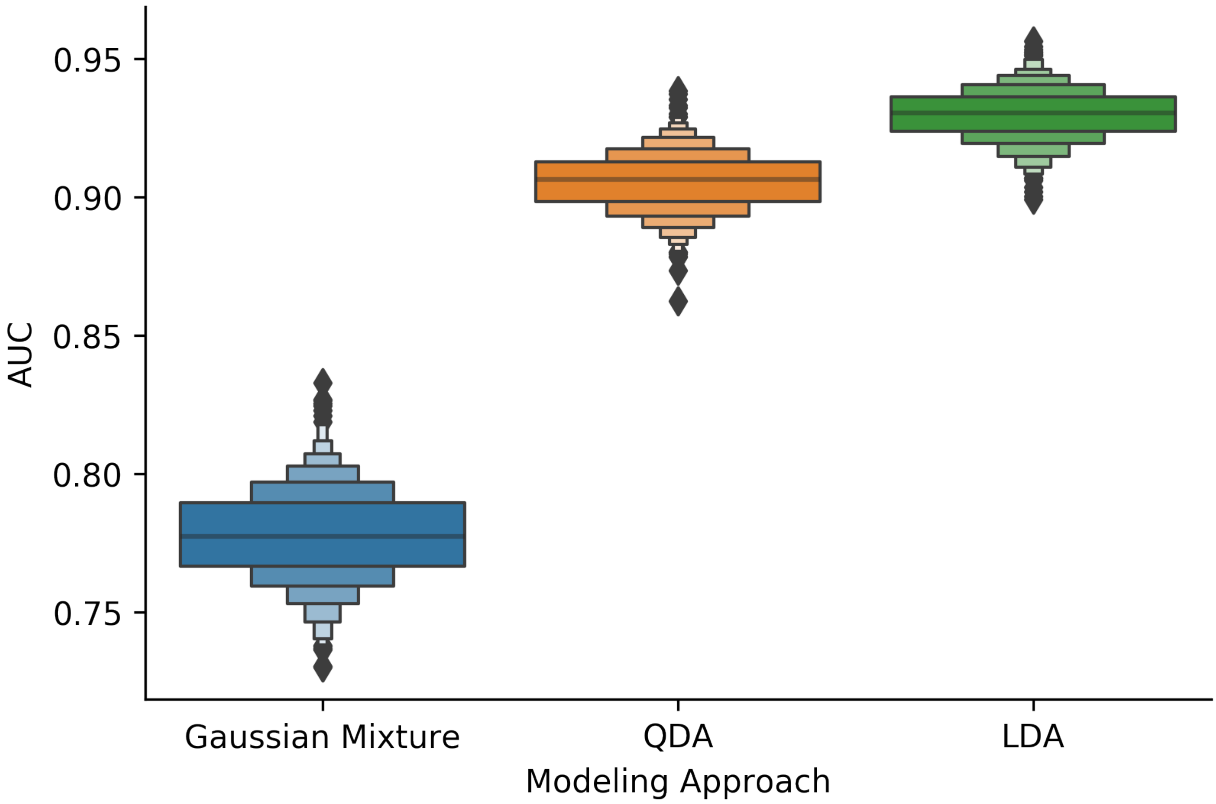


**Supplementary Table 2:** Diagnostic accuracy of Gaussian Mixture Model, Quadratic Discriminant Analysis, and Linear Discriminant Analysis models. Results are reported for 5-Fold cross-validated area-under-the-curve (AUC) estimates bootstrapped for each modeling approach.

| ***Approach*** | ***Median*** | ***2.5%*** | ***97.5%*** |
|---|---|---|---|
| ***Gaussian Mixture*** | 0.78 | 0.74 | 0.81 |
| ***QDA*** | 0.91 | 0.88 | 0.93 |
| ***LDA*** | 0.93 | 0.91 | 0.95 |

### ASSIGNMENT OF ABNORMAL PROBABILITIES FOR AMBIGUOUS TESTS

**Supplementary Figure 7**: Kernel density estimates of distribution of predicted abnormal probabilities based on whether maneuver was: (A) normal, (B) ambiguous, and (C) abnormal; these estimates were performed for the Deep Learning (DL), Traditional Logistic Regression (LR) and Hybrid approaches

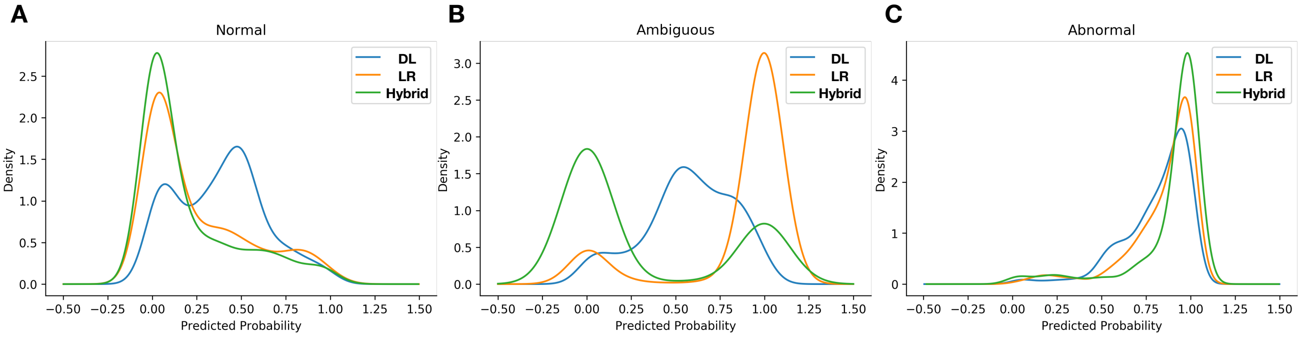


**Supplementary Table 3**: Odds ratios reported via univariable logistic regression that a predicted probability closer to 0.5 is related to the presence of an abnormal test; odds ratio greater than one indicates positive association of intermediate abnormal probability being association with abnormal test

| *Approach* | *Odds Ratio* | *CI[2.5%]* | *CI[97.5%]* | *P-Value* |
|---|---|---|---|---|
| *Deep Learning* | 4.21 | 2.78 | 6.38 | 1.16E-11 |
| *Traditional* | 1.07E-06 | 6.16E-08 | 1.84E-05 | 3.25E-21 |
| *Hybrid* | 2.72E-05 | 1.50E-06 | 4.95E-04 | 1.22E-12 |

**Supplementary Table 4:** Average predicted probabilities based on whether maneuver was ambiguous, normal, or abnormal

| *Approach* | *Ambiguous* | *Normal* | *Abnormal* |
|---|---|---|---|
| *Deep Learning* | 0.57 | 0.38 | 0.81 |
| *Traditional* | 0.86 | 0.25 | 0.85 |
| *Hybrid* | 0.31 | 0.20 | 0.89 |

### PROJECTION OF COMMON PREDICTORS BETWEEN TRADITIONAL AND DEEP LEARNING APPROACHES VIA CANONICAL CORRELATION ANALYSIS

**Supplementary Figure 8**: Embedding plots from canonical correlation analysis, with: (A) projection of traditional predictors towards a shared set of latent predictors between traditional/deep learning predictors; (B) projection of deep learning predictors based on common predictors

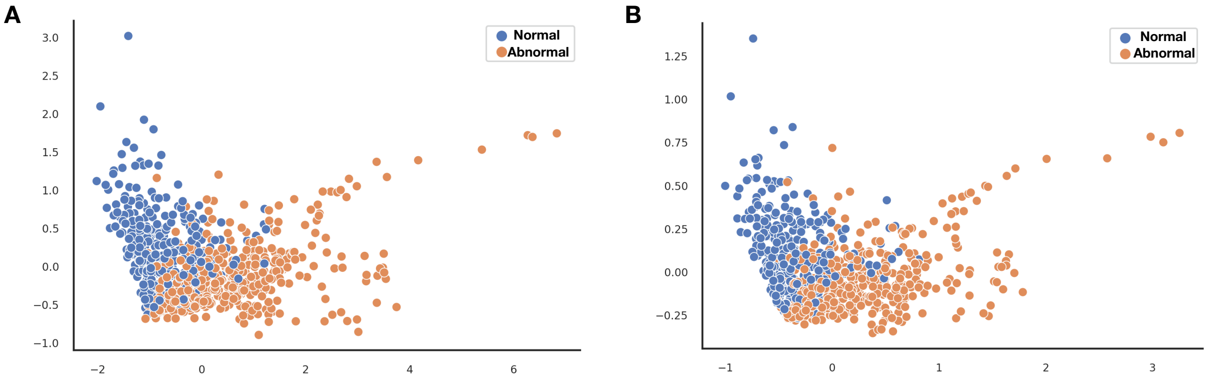

### IDENTIFYING COMPLEMENTARY PREDICTORS THROUGH SHAP PLOTS

**Supplemental Figure 9**: SHAP plots rank top important predictors (from most to least informative) based on predictors from: a) traditional approach, b) hybrid approach. Some deep learning predictors were found to be more important than the traditional predictors. Some of these predictors (eg. *DL Pred 48*, Deep Learning Predictor 48) were outside of common features identified by the CCA analysis, which suggests that deep learning approaches offer additional nuances not found by manual predictor extraction that may prove useful on more difficult use cases.

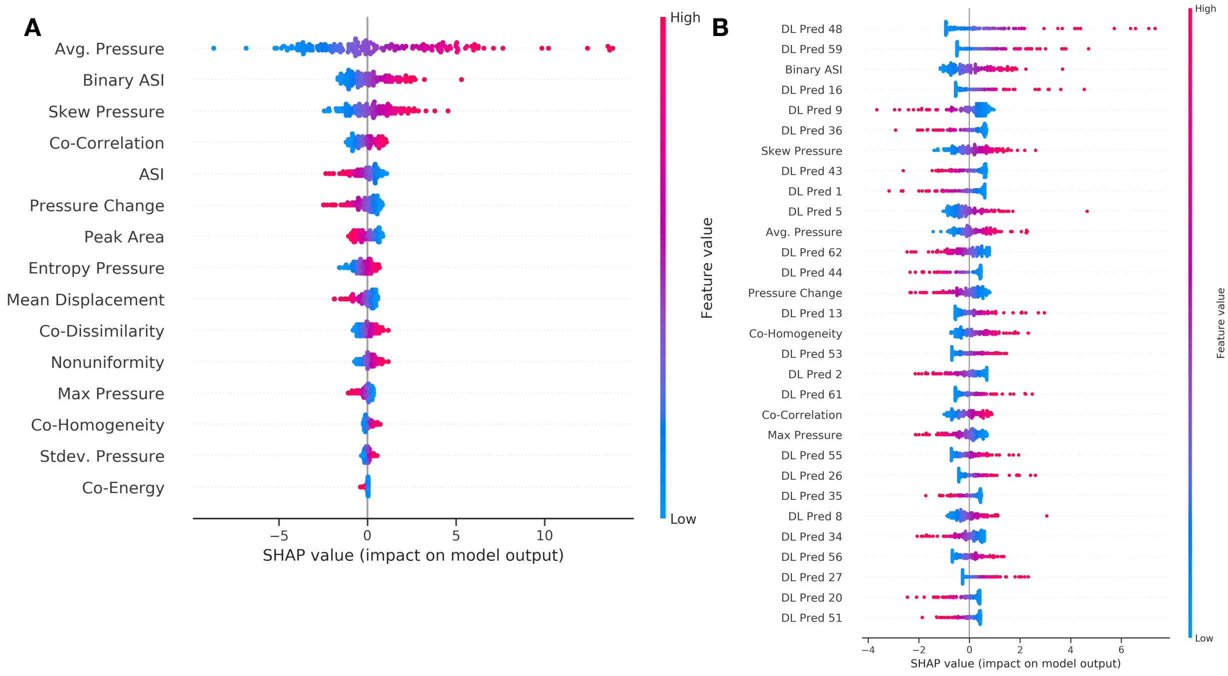
